# In vitro activity evaluation of *Lippia alba* essential oil against Zika virus

**DOI:** 10.1101/2020.06.25.170720

**Authors:** Bernardo E. Quispe-Bravo, Lucas Augusto Sevilla Drozdek, Joe Hermosilla Jara, Ingrit Elida Collantes Díaz, Edison Luiz Durigon, Enrique Walter Mamani Zapana, Egma Marcelina Mayta Huatuco, Juan Samuel Sulca Herencia

**Author notes:** Centro de Investigación de Genética y Biología Molecular, Universidad de San Martin de Porres, Lima, Peru (Bernardo E. Quispe-Bravo). These authors contributed equally to his work (Bernardo E. Quispe-Bravo, Lucas Augusto Sevilla Drozdek, Joe Hermosilla Jara, Juan Samuel Sulca Herencia). These authors also contributed equally to this work (Ingrit Elida Collantes Díaz, Edison Luiz Durigon, Enrique Walter Mamani Zapana, Egma Marcelina Mayta Huatuco).

## Abstract

III.

The Zika virus (ZIKV) of the *Flaviviridae* family is an emerging virus that caused, between 2016 and 2018, serious public health problems in Latin America, affecting neonates with greater severity. The clinical spectrum includes Guillain-Barré syndrome, microcephaly and others neurodegenerative diseases. There is no antiviral treatment or vaccine against this virus, for that reason the antiviral properties of various plants are being studied. *Lippia alba*, locally known as “Prontoalivio”, is an aromatic shrub of the *Verbenaceae* family with a wide geographical distribution (especially in South and Central America) and is used in traditional medicine against fever, skin diseases and as a pain reliever. In this study, the antiviral activity of the essential oil of *Lippia alba* against ZIKV was evaluated in the Vero 76 cell line. *Lippia alba* was collected in the department of Amazonas, in the rainforest of Peru, and identified in the Museo de Historia Natural of the Universidad Nacional Mayor de San Marcos. The essential oil sample was obtained by steam hydrodistillation. The essential oil showed cytotoxicity to a concentration greater than or equal to 167 μg/mL in the Vero 76 cell line. The antiviral activity of essential oil against ZIKV (previously identified by real-time PCR and propagated in the C6/36 cell line) was evaluated using the plaque reduction test (PRP). The essential oil showed antiviral activity in concentrations from 8.02 μg/mL to 20.88 μg/mL, which represents a range between 59.44% to 85.56% of plaque reduction and may be considered as a candidate for antiviral studies against ZIKV.

**Author Summary:** Environmental temperature fluctuations, human activities and vector characteristics increase ZIKV cases worldwide. This neglected disease has been silently affecting people of all ages, generating greater impact on neonates by causing microcephaly and other neurodegenerative diseases. ZIKV vaccines are in phase II trials and there is no antiviral treatment. Nowadays, the study of antiviral plants is gaining strength in the scientific community because they are known to contain chemical compounds that could be drugs candidates. In Peru, there are no antiviral treatments studies reported against Zika virus, this first report is important because it creates a new line of research for future studies of antiviral plant extracts against neglected viral-diseases. Finally, the essential oil of *Lippia alba* showed antiviral activity against ZIKV in Vero 76 cells, moreover, we intend to carefully isolate and study the chemical compounds as drug candidates in animals trials and possible humans trials.

**Disclaimer:** The views expressed in this manuscript are those of the authors and do not necessarily reflect the official policy or position of the Department of the Navy, Department of Defense, nor the U.S. Government.

**Copyright statement:** Juan Sulca is an employee of the U.S. Government. This work was prepared as part of their official duties. Title 17 U.S.C. § 105 provides that ‘Copyright protection under this title is not available for any work of the United States Government’. Title 17 U.S.C. § 101 defines a U.S. Government work as a work prepared by a military service member or employee of the U.S. Government as part of that person’s official duties.

## V. Introduction

In 2019, the World Health Organization (WHO) reported that 87 countries have evidence of autochthonous transmission of Zika virus (ZIKV) (1). In 2016, the first case of autochthonous transmission was reported in Peru, where *Aedes aegypti* is the main vector of this disease (2). ZIKV disease presents fever, malaise, headaches, rash, conjunctivitis, muscle and joint pains, and neurological complications such as neuropathies, myelitis and Guillain-Barré syndrome (GBS) (1,2). Pregnant women infected with this disease could present GBS and adverse pregnancy outcomes such as premature delivery, fetal death and congenital malformations (1,2). Currently, there are no antiviral treatments or vaccines against this disease, so the study of medicinal plants is of great importance because they could help us identify new chemical compounds with antiviral properties. These compounds can be subsequently evaluated using in silico approaches to study their interactions with possible viral protein targets. These compounds can also be used in pre-clinical evaluations in animal models and in human clinical trials (3). Within the *Verbenaceae* family, the genus *Lippia* contains about 220 species, *Lippia alba* is a shrub widely distributed in different regions of Peru (4,5) and presents various ethnopharmacological applications to relieve problems such as digestive, respiratory, cardiovascular, anemia, headache and others (6). It has also been proven to have antiviral activity against viruses such as influenza virus type A (IV A), poliovirus type 2 (PV 2) and flaviviruses such as yellow fever virus (YFV) and dengue virus serotype 2 (DENV 2), which are closely related to ZIKV (7–10). However, no published studies evaluating its in vitro antiviral potential against ZIKV were found. Therefore, the present study aims to investigate the antiviral activity of *Lippia alba* against ZIKV in the Vero 76 cell line.

## VI. Methods

### Online Protocol

DOI: dx.doi.org/10.17504/protocols.io.bhpyj5pw

HTML: https://protocols.io/view/prt-assay-for-antiviral-activity-of-the-essential-bhpyj5pw.html

### Method

1. WORK EXECUTION The investigation was carried out at the Laboratorio de Virología Clínica y Molecular of the Universidad Nacional Mayor de San Marcos in Lima, Peru.
2. HARVEST AND IDENTIFICATION OF *Lippia alba* The shrub was collected in the community of Miraflores, Bagua Grande, Amazonas, Peru (S 5°52,534’; W 78°23.087’; elevation of 1,699 MASL). The species was confirmed as *Lippia alba* at the Museo de Historia Natural of the Universidad Nacional Mayor de San Marcos (Registration number: 300373).
3. CELL LINE AND VIRAL STRAIN The ZIKV viral strain, the C6/36 (ATCC CRL 1660) and Vero 76 (ATCC CRL 1587) cell lines were donated by the Laboratorio de Virología Clínica y Molecular of the Universidade de São Paulo.
4. ESSENTIAL OIL EXTRACTION OF *Lippia alba* The essential oil was extracted from the entire plant using a steam-driven hydrodistillation system (11). The plant sample was minced and deposited into borosilicate glass round-bottomed flask with short neck and then covered with bi-distilled water (11). The hydrodistillation process was carried out for 3 hours (12) using petroleum ether as the extraction solvent (13) and then dried over solid anhydrous sodium sulfate (13,14). The petroleum ether residues were evaporated at room temperature and stored at 4 °C in darkness (13,14).
5. CELL LINE PROPAGATION: ZIKV INFECTION AND HARVEST The methodology of the Instituto de Medicina Tropical “Pedro Kouri” (IPK) was used with some modifications (15). C6/36 and Vero 76 cell lines were seeded in medium at pH 6.9 in 6-well plates at a density of 1.25 × 10^5^ cells/mL and 2.5 × 10^5^ cells/mL, respectively. The C6/36 and Vero 76 cell lines are incubated at 28°C and 37°C, respectively, (2mL per well) 2 days before infection. Cell monolayer growth was confirmed 2 days after seeding. The cell infection was carried out inoculating 100 μL/well of ZIKV, incubated for 1 hour and then maintenance medium was added. It was incubated until the appearance of the cytopathic effect (CPE). Then the supernatant was centrifuged at 300 rpm/10min/4°C, aliquoted and stored at −80°C. Viral passages were performed until the viral seed shows an optimal cytopathic effect in Vero 76 cells.
6. VIRAL TITER A ten-fold serial dilution of ZIKV with maintenance medium (MM) was carried out and incubated at 37°C for 1 hour. Immediately, 50 μL/well of ZIKV was inoculated in triplicate for each dilution in 24-well plates with confluent Vero 76 cell monolayer and incubated at 37°C for 3 hours. Then, 500 μL/well of overlay medium containing carboxymethyl cellulose was added and incubated at 37°C for 6 days. The assay was stained with naphthol blue black (NBB) and the viral titer was obtained by counting the number of plaque-forming units per milliliter (PFU/mL).
7. CYTOTOXICITY TEST Tween 80 and ethanol were mixed in a proportion of 2:1 and then mixed with the essential oil of *Lippia alba* (12) forming the essential oil mixture (EOM) or stock solution. The EOM dilutions were carried out with maintenance medium (MM) and incubated at 37°C for 1 hour. Finally, 125 μL/well of each EOM dilution was inoculated in triplicated into a 96-well plate with Vero 76 cells in a confluent monolayer and incubated at 37°C. The assay was stained on day 6 with NBB. The mean cytotoxic concentration (CC_50_) is defined as the minimum dilution of the compound that induces the death of 50% of the Vero 76 cells and the calculation is expressed in μg/mL (16).
8. PLAQUE REDUCTION TEST (PRT_50_) The EOM dilutions without cytotoxity were mixed in equal parts with the viral solution and incubated at 37°C for 1 hour (the dilutions of the essential oil have a final concentration between 0.65 μg/mL to 167 μg/mL). Subsequently, 50 μL/well of each new dilution was inoculated in triplicate into a 24-well plate with Vero 76 cells in a confluent monolayer and incubated at 37°C for 3 hours. Finally, the overlay medium containing carboxymethyl cellulose was added and incubated at 37°C for 6 days, then assay was stained with NBB. A virus and cell controls were used and the essential oil of EOM was replaced with the maintenance medium to control for the antiviral activity of Tween 80 and ethanol against ZIKV. We considered that there was a positive antiviral activity if the mix inhibited the formation of 50% of the inoculated plaque-forming units in well.

## VII. Results

The ZIKV harvests (fourth viral passage) in the C6/36 and Vero 76 cell lines had a viral titer of 9.8×10^6^ PFU/mL and 2.6×10^4^ PFU/mL, respectively. So, it was decided to carry out the trials with C6/36 cell line.

Total cell death (100% cytotoxicity) in the Vero 76 cell line was obtained with concentrations greater than 668 μg/mL of the essential oil of *Lippia alba*, with a CC_50_ of 334 μg/mL (Table1). The concentrations equal to or less than 167 μg/mL did not show a cytotoxic effect and are optimal for Plaque Reduction Test (PRT).

**Table 1:** Cytotoxic evaluation of the essential oil of *Lippia alba* in the Vero 76 cell line.

| Extract<br>concentration<br>(µg/mL) | EOM sample |  |  | Control |  |
| --- | --- | --- | --- | --- | --- |
|  | Cells with<br>MM<br>(%) | Tween 80 and ethanol<br>+<br>Oil essential |  | Tween 80 and ethanol<br>+<br>MM |  |
|  |  | CC | CC (%) | CC | CC (%) |
| 4008 | 100 | + | 100 | + | 100 |
| 1336 | 100 | + | 100 | + | 100 |
| 668 | 100 | + | 100 | + | 100 |
| 400.8 | 100 | + | 75 | - | 0 |
| 334 | 100 | + | 50 | - | 0 |
| 250.5 | 100 | + | 25 | - | 0 |
| 167 | 100 | - | 0 | - | 0 |
| 83.5 | 100 | - | 0 | - | 0 |
| 41.75 | 100 | - | 0 | - | 0 |
| 20.88 | 100 | - | 0 | - | 0 |
| 13.36 | 100 | - | 0 | - | 0 |
| 10.44 | 100 | - | 0 | - | 0 |
| 8.02 | 100 | - | 0 | - | 0 |
| 5.22 | 100 | - | 0 | - | 0 |

CC: Cytotoxicity effect expressed qualitatively through presence (+) or total-absence (-) of cell-death.
CC(%): Cytotoxic effect expressed as a percentage of the proportion of dead cells in the cellular monolayer.
“+” There is a cytotoxic effect.
“-“ There is no cytotoxicity .

Using PRT_50_, it was shown that the essential oil of *Lippia alba* inhibited ZIKV at concentrations in the range of 8.02 to 20.88 μg/mL. The concentrations greater than 20.88 μg/mL could not be evaluated due to the presence of high concentrations of Tween 80 and ethanol, as their mixture alone caused inhibition of the ZIKV (Table 2)

**Table 2:** Evaluation of the antiviral activity of *Lippia alba* essential oil against ZIKV using of PRT_50_ in the Vero 76 cell line.

| Essential oil concentration (µg/mL) | PFU average of ZIKV | EOM sample |  | Control |  |
| --- | --- | --- | --- | --- | --- |
|  |  | Tween 80 and ethanol + Essential oil |  | Tween 80 and ethanol + Maintenance medium |  |
|  |  | PFU/well | % Reduction | PFU/well | % Reduction |
| 167 | 18 | 0 | 100 | 0 | 100 |
| 83.5 | 18 | 0 | 100 | 0 | 100 |
| 41.75 | 18 | 0 | 100 | 0 | 100 |
| 20.88 | 18 | 2.6 | 85.56 | 12.85 | 28.61 |
| 13.36 | 18 | 3.7 | 79.44* | 18 | 0 |
| 10.44 | 18 | 4.9 | 72.78 | 18 | 0 |
| 8.02 | 18 | 7.3 | 59.44 | 18 | 0 |
| 5.22 | 18 | 9.3 | 48.33* | 18 | 0 |
| 4 | 18 | 18* | 0 | 18 | 0 |
| 2.61 | 18 | 18 | 0 | 18 | 0 |
| 1.3 | 18 | 18 | 0 | 18 | 0 |
| 0.65 | 18 | 18 | 0 | 18 | 0 |

## VIII. Discussion

Firstly, our results showed that it is possible to propagate and harvest ZIKV using the C6/36 cell line (isolated from salivary glands of *Aedes albopictus*), in less time and with higher viral titers. This result was established on the basis of a comparison between C6/36 and Vero 76 cell lines. Other studies have used the C6/36 cell line as a model for the viral propagation of dengue, yellow fever and chikungunya viruses (17). Mosquito cell line (the C6/36 cell line included) have a persistent non-lethal infection due to their replication-limiting signaling pathways, such as: the Janus kinase/signal transducers and activators of transcription (JAK-STAT), Toll-like receptors (TLRs) and RNA interference (RNAi) (18). Mosquitoes have a low internal temperature compared to mammals, which slows down viral replication kinetics (19) and maintains the stability of the 3’-terminal stem loop (3’SL) expressing a persistent non-pathogenic character in flavivirus infections (18). Alternating hosts and high viral passages could generate mutations and nucleotide variations due cell-receptors exchange with binding viral-protein (20,21). Therefore, Zika viral fourth-passage was chosen in the C6/36 cell in order to reduce or avoid genetic changes, moreover, this passage was chosen because it produces well-defined lytic plaques using 3% carboxymethyl cellulose (as a semi-solid overlay medium). This method is widely used in other plaque reduction tests for flavivirus. For example, the antiviral activity of sofosbuvir against the yellow fever virus (22), as well as some fungal chemicals against DENV2 (23).

The essential oil of *Lippia alba* was shown to reach a CC_50_ in the Vero 76 cell line at a concentration of 334 μg/mL, other similar studies reported lower values (in the range of 12.3 to >200 μg/mL) (8,9,24). No assays above 668 μg/mL were performed because the high concentrations of the Tween 80 and ethanol mixture, which were shown to generate a total cytotoxicity in Vero 76. Moreover, essential oil mixture (conformed by Tween 80, ethanol and essential oil) showed partial-cytotoxicity in Vero 76 cells in the range of 250.5 to 400.8 μg/mL. The differences in the cytotoxic activity of *Lippia alba* in Vero 76 cells could be directly related to the different concentrations of the components of the essential oil which varies due to environmental factors, harvest area and methodology for separacting the essential oil from the solvent used (25,26). Moreover, each component of the essential oil of *Lippia alba*, including beta-caryophyllene, nerol, Geraniol, Citronellol, Eugenol, Citral and Carvone have different cytotoxic effects in Vero 76 cells at different concentrations: 39.7, 47.8, 59.3, 64.4, 71.3, 124.1 and >200 μg/mL, respectively (16).

The essential oil of *Lippia alba* reduced lytic plaque formation by more than 50% using concentrations between 8.02 and 20.88 μg/mL. In similar studies, which evaluated the same essential oil and its cytotoxic activity against other flaviviruses, reported that the total essential oil of *Lippia alba* inhibits DENV 2 at concentrations between 0.4 and 32.6 μg/mL (9) and against YFV in a range between 4.3 and 25 μg/mL (27). The essential oil concentrations values above 20.88 μg/mL could not be evaluated because the EOM sample contains a high concentration of Tween 80 (emulsifier) and ethanol (solvent) which reduce the lytic plaque-forming of Zika virus. Tween 80 is a non-ionic detergent with very low micellar concentration, moreover, it is hydrolyzed by cellular enzymes such as lipases and esterases, producing oleic acid (fatty acids) which is a powerful viricidal agent. The oleic acid, which contains 18 carbon chains, binds to the viral envelope protein through strong association due to the high number of carbon chains. Moreover, oleic acid is attaches to viral envelope proteins and the draws them out of the viral surface, forming protein-lipid complexes, avoiding the entry of the viral genome inside the host by adsorption (28,29). Additionally, high temperatures generate faster virus inactivation due to accelerated hydrolysis of PS80 and subsequent interactions between oleic acid and the viral envelope (29). With respect to the solvent, ethanol induces the activity of the enzyme 2’,5’ oligoadenylate synthetase whose principal role is to promote cellular resistance against viral infection (developing an antiviral state). Furthermore, it has been shown that the ethanol induces the production of interferon-beta (IFN-β) which is secreted by fibroblasts in response to viral stimuli. Interferon-beta is secreted and released extracellularly, binds to its cell membrane receptor and enters the cell (30,31). The presence of interferon-beta (ethanol-induced) induce the 2’,5’ oligoadenylate synthetase activity and antiviral activity (31). The enzyme, in the presence of viral mRNA, activates endoribonucleases that degrade and inhibit the translation of viral mRNA, preventing viral replication and promoting antiviral activity (30,31). The essential oil of *Lippia alba* and culture medium are immiscible, so a surfactant or emulsifier (Tween 80) to homogenize the mixture and a solvent (ethanol) to decrease viscosity were used. Tween 80 inactivates viruses, and has a low micellar concentration. Other emulsifiers are similar but more toxic, like Triton X-100 (29).

*Lippia alba* has been shown to possess antiviral activity against Zika virus and its chemical components have been shown to have broad-spectrum antiviral properties against flaviviruses and some other viruses tested in other studies, including DENV 2, IV A, PV 2 and YFV (7–10,27). So, due to the great variety of chemical components in the essential oil from *Lippia alba*, it is important to carry out in silico screening to demonstrate the synergy or non-synergy of the chemical components against ZIKV (16,32). Because a study carried out in 2016, where the antiviral activity of H. *cordata* against DENV 2 was evaluated, reporting that quercetin and quercetrin have low antiviral activity, however, the synergy of these components enhanced their antiviral activity against DENV 2 (33).

Molecular docking predicts the high-affinity binding sites of a protein-ligand complex using three-dimensional structure through in silico assays (34). Molecular docking analysis of the chemical compounds found in the essential oil to *Lippia alba* against non-structural and structural-proteins of ZIKV has shown a high affinity of coupling with geraniol acetate, bicyclosesquiphellandrene, α-copaene, β-bourbonene, geraniol, geranial, germacrene and caryophyllene oxide, being these possible candidates drugs against ZIKV (35–37). In vitro, beta-caryophyllene has antiviral activity against the non-structural protein NS1 of DENV 2 (38) and citral has antiviral activity against YFV and DENV 2 (27,38). Caryophyllene and citral could present antiviral activity against ZIKV (before or after the adsorption) because it also belongs to the same flavivirus-family.

Finally, through in vitro trials, the unfractionated essential oil produced by steam-hydrodistillation of the *Lippia alba* demonstrated antiviral activity against ZIKV. Moreover, we recommend analyzing and characterizing the essential oil of *Lippia alba* employing phytochemical screening and in silico assays, such as molecular docking, to predict which components could have the strongest antiviral activity as stand-alone or in synergy with other compounds for future in vitro and in vivo assays.

## IX. Acknowledgments

We are grateful to PhD. Claudio N. Villegas-Llerena for his thoughtful comments during the translation of this manuscript.

